# Quantitative modeling demonstrates format-invariant representations of mathematical problems in the brain

**DOI:** 10.1101/2022.04.18.488703

**Authors:** Tomoya Nakai, Shinji Nishimoto

## Abstract

Mathematical problems can be described in either symbolic form or natural language. Previous studies have reported that activation overlaps exist for these two types of mathematical problems, but it is unclear whether they are based on similar brain representations. Furthermore, quantitative modeling of mathematical problem solving has yet to be attempted. In the present study, subjects underwent 3 h of functional magnetic resonance experiments involving math word and math expression problems, and a read word condition without any calculations was used as a control. To evaluate the brain representations of mathematical problems quantitatively, we constructed voxel-wise encoding models. Both intra- and cross-format encoding modeling significantly predicted brain activity predominantly in the left intraparietal sulcus (IPS), even after subtraction of the control condition. Representational similarity analysis and principal component analysis revealed that mathematical problems with different formats had similar cortical organization in the IPS. These findings support the idea that mathematical problems are represented in the brain in a format-invariant manner.

## Introduction

Our mathematical problems can be presented in different formats. In symbolic form [math expression (ME)], mathematical problems are represented as a combination of digits and operators (e.g., “3 × 2 = 6”); in linguistic form [math word (MW)], the same mathematical problems are described in natural language (e.g., “There are 3 eggs in each box. How many eggs are in 2 boxes?”). Although most mathematical problems are represented in symbolic ME format, the MW format is still “one of the most common materials in school curricula for teaching students to transfer mathematical knowledge into real-world contexts” (Ng *et al*., 2021). Studies with a large sample of schoolchildren have found that performance in MW problems is related to later performance in more advanced algebra (Fuchs *et al*., 2012; Powell & Fuchs, 2014), suggesting that a shared cognitive basis of mathematical problem solving may exist across different formats.

In several neuroimaging studies, the format-independency of mathematical problem solving has been examined using univariate methods (Sohn *et al*., 2004; Newman *et al*., 2011; Zhou *et al*., 2018; Chang *et al*., 2019). Sohn *et al*. reported that MW problems induce activity in the left prefrontal cortex, whereas ME problems induce activity in the bilateral parietal cortex (Sohn *et al*., 2004). Newman *et al*. found evidence for shared activation of MW and ME problems in the bilateral intraparietal sulcus (IPS) and left frontal cortex (Newman *et al*., 2011). Zhou *et al*. tested MW problems with different complexities and found activations that included the left inferior frontal gyrus and dorsomedial prefrontal cortex (Zhou *et al*., 2018). Chang *et al*. found that, compared with sentence comprehension, MW problems induced higher activation in the IPS and dorsolateral prefrontal cortex (Chang *et al*., 2019). Collectively, these studies suggest the involvement of the IPS and prefrontal cortex in both MW and ME problem solving. However, it remains unclear whether the two types of mathematical problems share brain representations. Furthermore, quantitative modeling of mathematical problem solving has yet to be attempted.

To address these issues, we used a voxel-wise encoding modeling approach (Naselaris *et al*., 2011). Encoding models predict brain activity in a quantitative manner based on a combination of features extracted from the presented stimuli. Researchers have adopted this approach to examine brain responses to visual (Kay, Naselaris, *et al*., 2008; Nishimoto *et al*.,2011; Huth *et al*., 2012; Çukur, Huth, *et al*., 2013; Çukur, Nishimoto, *et al*., 2013), auditory (De Angelis *et al*., 2018; Nakai, Koide-Majima, *et al*., 2021), semantic (Huth *et al*., 2016; Kiremitçi *et al*., 2021; Nakai, Yamaguchi, *et al*., 2021; Nishida *et al*., 2021; Shahdloo *et al*., 2022), and emotional (Horikawa *et al*., 2020; Koide-Majima *et al*., 2020) stimuli, and to assess many complex cognitive functions (Nakai & Nishimoto, 2020). In addition, the approach allows for quantitative evaluation of the generalizability of models constructed under a certain format to data in another format (Nakai, Yamaguchi, *et al*., 2021). Moreover, encoding models enable visualization of multivoxel patterns with different categorical information and evaluation of their similarity across different formats.

In the current study, subjects underwent 3-h functional magnetic resonance imaging (fMRI) experiments (**Figure 1A**) and each attempted to solve a series of mathematical problems in the MW and ME formats. As a control for non-math features such as visual attention, motor control, and linguistic processing, we also included a read word (RW) condition, under which the subjects read the same sentences as under the MW condition but did not conduct calculations (see Methods for details). We prepared the control condition only for the MW condition as this condition would be more demanding than a control corresponding to the ME condition, thus suitable for eliminating a possible influence by features of noninterest. Using sparse operator features, we modeled brain activity specific to each mathematical problem (**Figure 1B**). By applying the cross-modal modeling technique developed in our previous study (Nakai, Yamaguchi, *et al*., 2021), we defined a format invariance (FI) index, which quantifies how much information can be captured in a format-invariant manner in each brain region. In addition, representational similarity analysis (RSA) and principal component analysis (PCA) revealed representational relationships among different mathematical problems. Thus, through quantitative evaluations, this study provides new evidence of brain representations in math problem solving.

**Figure 1.**
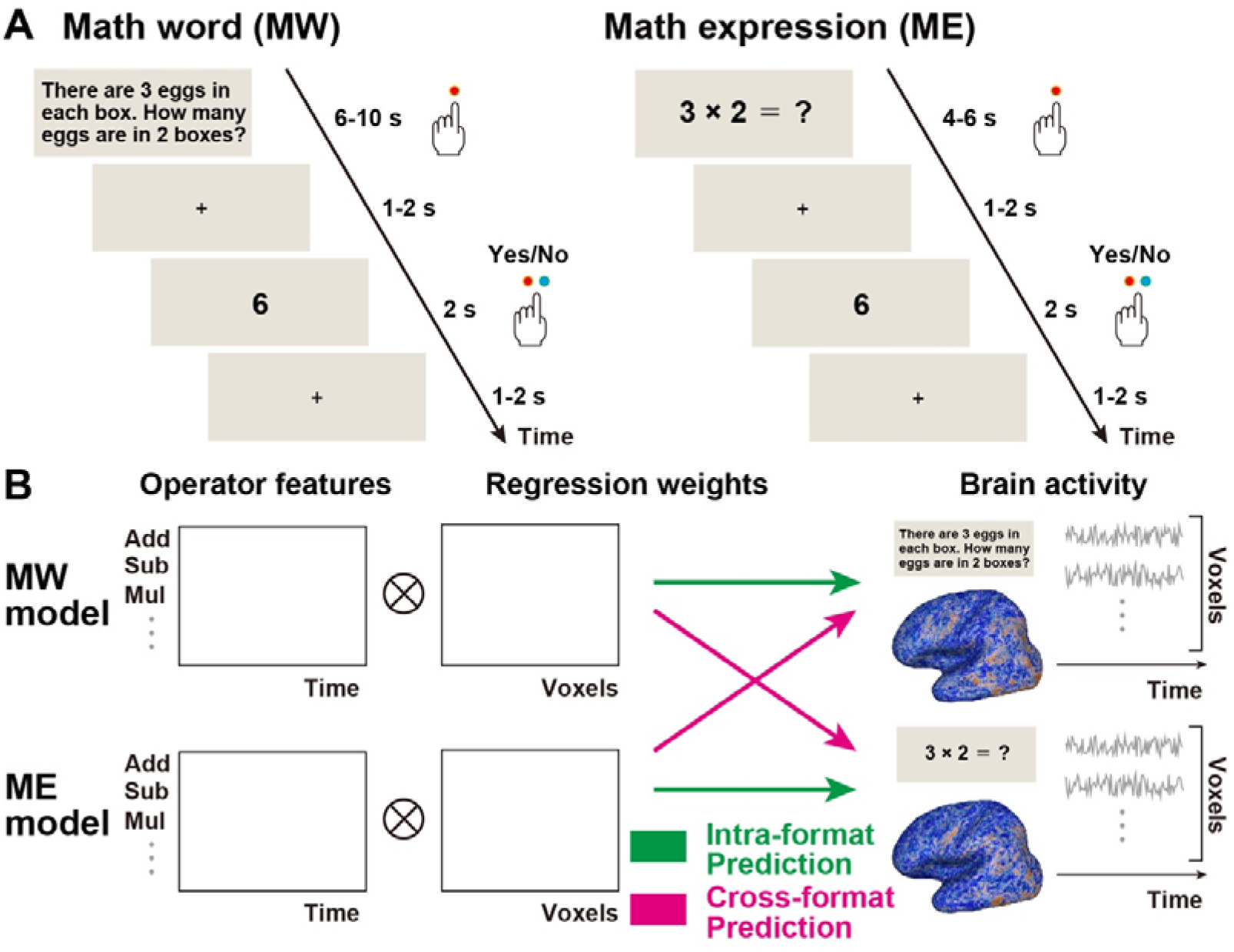
Experimental design. (**A**) In the math word (MW) and math expression (ME) conditions, mathematical problems were presented either in a natural language format for 6–10 s or in a symbolic format for 4–6 s. Subjects pressed the button on the left when they had solved a given problem. They pressed the button on the left or right to indicate whether the probe digit stimulus (presented for 2 s) did or did not match their answer, respectively. The subjects pressed the button on the left when they remembered all nouns that appeared in the given sentence. The probe stimuli consisted of two nouns; the subjects pressed the buttons on the left or right to indicate whether both nouns were or were not included in the previous sentences, respectively. (**B**) Intra- and cross-format encoding models were constructed to predict brain activity from operator features. The feature matrix and brain response matrix are transposed for the purpose of visualization.

## Results

### Encoding models predict brain activity in an intra- and a cross-format manner for each subject

To confirm that the encoding models successfully captured brain activity during the math problem solving, we performed a series of intra-format encoding modeling predictions and examined the prediction accuracy using a test dataset with the same format as that of the training dataset (**Figure 1B**, green). First, we trained an encoding model using the MW training dataset and predicted brain activity using the MW test dataset. Using operator features, the encoding model significantly predicted the activity in large brain networks, including the bilateral inferior frontal, parietal, and occipital cortices *(P* < 0.05, FDR corrected; 40.1% ± 8.2% of voxels were significant; **Figure 2A** blue; **Table S1**). Similarly, we trained encoding models using the ME training dataset and predicted brain activity using the ME test dataset. This encoding model significantly predicted a smaller part of the region, including the bilateral inferior frontal, bilateral parietal, and bilateral occipital cortices, compared with that predicted by the MW intra-format encoding model (21.5% ± 9.4% of voxels were significant; **Figure 2B** blue; **Tables S1**).

**Figure 2.**
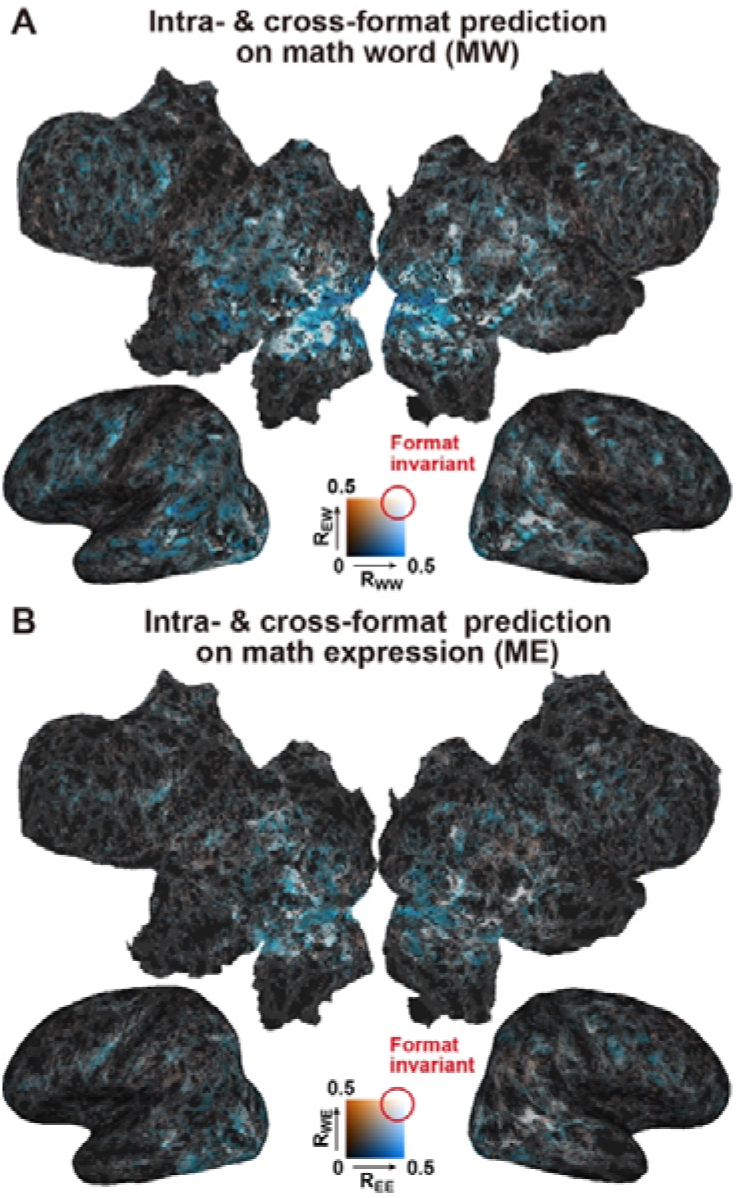
Intra- and cross-format predictions. (**A**) Comparison of the prediction accuracies when applied to the math word (MW) test dataset using operator features. The intra-format prediction accuracies of the MW model (denoted as R_WW_, blue) and the cross-format math expression (ME) model (denoted as R_EW_, orange) are mapped onto the cortical surface of subject ID01. Regions in which activity was predicted, regardless of the stimulus format, are shown in white. Only significant voxels *(P* < 0.05, FDR corrected) are shown. (**B**) Comparison between the prediction accuracies of the ME test dataset. Intra-format prediction accuracy, using the ME model (denoted as R_EE_, blue) and cross-format prediction accuracy, using the MW model (denoted as R_WE_, orange) are mapped onto the cortical surface.

To determine whether the encoding models were generalizable to mathematical problems in other formats, we performed cross-format encoding model analyses. Specifically, we applied an encoding model trained with the ME training dataset to the MW test dataset; prediction accuracy was significant in the bilateral inferior frontal, bilateral parietal, and bilateral occipital cortices (21.2% ± 8.2% of voxels were significant; colored in orange or white in **Figure 2A**; **Table S1**). Similarly, we applied the encoding model trained with the MW training dataset to the ME test dataset; prediction accuracy was also significant in the bilateral inferior frontal, bilateral parietal, and bilateral occipital cortices (12.2% ± 4.8% of voxels were significant; colored in orange or white in **Figure 2B**; **Table S1**). In summary, our encoding models successfully predicted brain activity induced in large brain regions, including the parietal and frontal cortices, across different formats of mathematical problems.

### Format-invariant and format-specific processing of mathematical problems

To quantify how much information a certain model captures for each format of mathematical problem, we calculated FI indices by combining the intra-format and cross-format prediction accuracies. FI was adopted based on modality invariance indices, which have been used previously to quantify modality-invariant processing of semantic information across text and speech modalities (Nakai, Yamaguchi, *et al*., 2021). We found clusters of significant FI values distributed across the bilateral frontal, parietal, and occipital cortices (19.7% ± 9.9% of voxels were significant across the whole cortex; **Figure S1**). To quantitatively evaluate the consistency of our results across test subjects, we calculated the ratio of significant voxels in each anatomical region of interest (ROI) of each subject and averaged across subjects. We focused on ROIs where >20% of voxels had a significant FI value. Suprathreshold FI values were found in large brain regions, including the bilateral prefrontal, parietal, and occipital cortices (**Figure 3A**).

**Figure 3.**
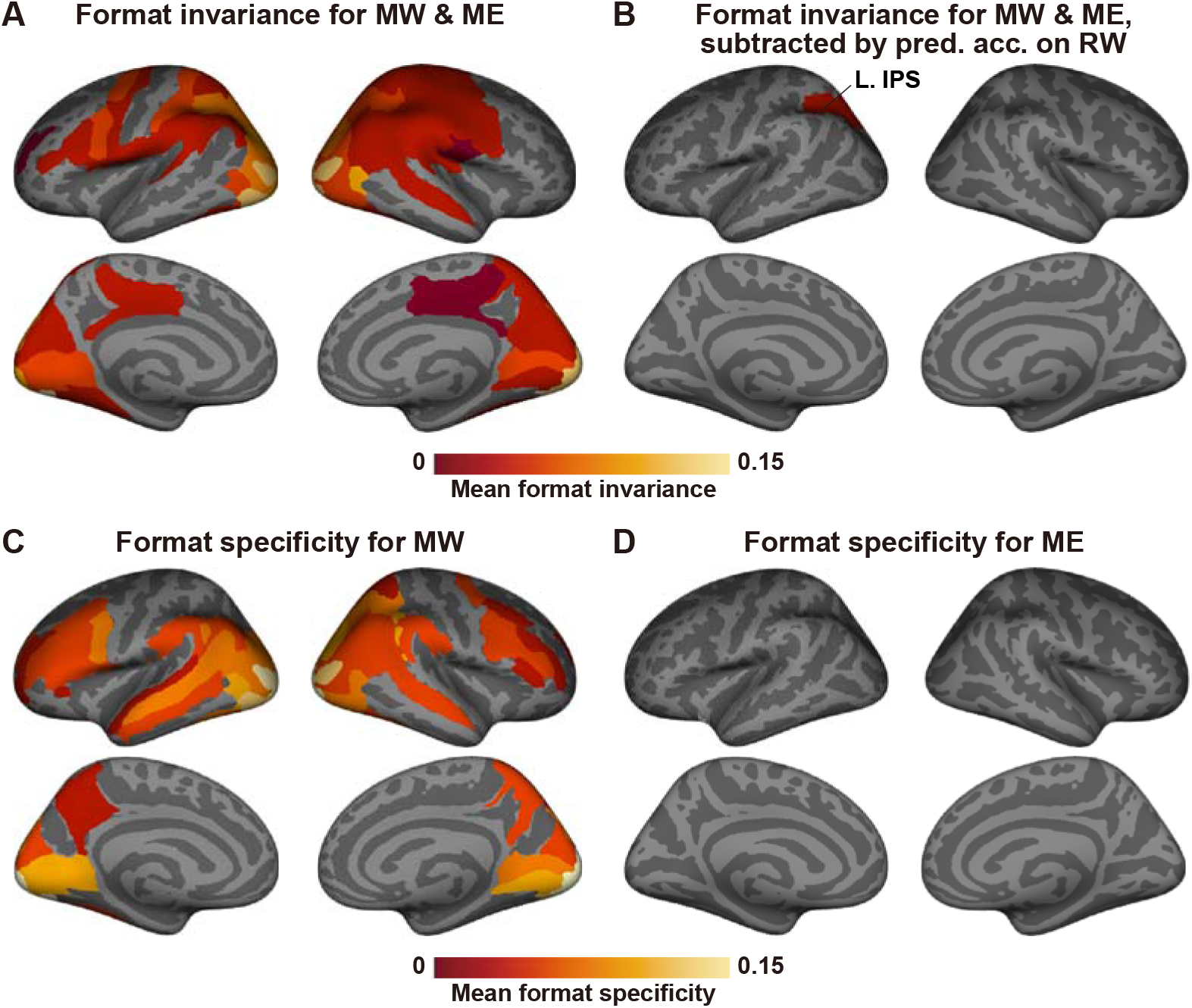
Format invariance and format specificity. (**A**) Group average of format invariance (FI) mapped onto the template brain. FI values were averaged in each anatomical ROI and averaged across eight subjects. Only ROIs with >20% significant voxels are shown. (**B**) Group average of FI with subtraction of the prediction accuracy in the read word (RW) condition. (**C, D**) Group average of format specificity (FS) for (**C**) MW and (**D**) ME, mapped onto the template brain. IPS, intraparietal sulcus; L, left hemisphere.

The original encoding models predict not only arithmetic-related regions but also regions related to sensorimotor and semantic information. To exclude regions of noninterest from our analyses, we applied encoding models trained in MW and ME conditions to the control RW condition. The average prediction accuracies of these two models were regarded as baseline accuracies. The baseline prediction accuracy was subtracted from the original intra- and cross-format prediction accuracies, and FI values were calculated. Consequently, we observed significant FI values predominantly in the left IPS (6.2% ± 3.8% of voxels were significant across the cortex; **Figure 3B**), indicating that format-invariant representations of mathematical problems are robust across different problem structures.

We used the RW condition (using the same visual stimuli as the MW condition) as a control for both MW and ME conditions because we assumed that the MW condition would induce larger activations than the ME condition. To confirm this assumption, we quantified format specificity (FS) values for both MW and ME conditions. FS was adopted based on modality specificity indices, which have been used previously to quantify modality-specific processing of semantic information for text or speech modalities (Nakai, Yamaguchi, *et al*.,2021). We found above threshold FS values only for MW condition (**Figure 3C**). In contrast, no above threshold FS values were found for the ME condition (**Figure 3D**), supporting our assumption that the RW condition serves as a control for both MW and ME conditions.

One concern regarding our analyses was whether the encoding model results could be explained solely by the problem complexity factor or also by cognitive load. To examine this further, we first constructed additional encoding models using only single - or double-operators (**Figure S2**); for both operator types, we found significant FI values in the bilateral IPS (10.4% ± 5.5% and 17.2% ± 8.1% of voxels were significant across the cortex, respectively), indicating that model predictability for the bilateral IPS was not a superficial effect due to differences between problem complexities. To further investigate our concern, we performed encoding modeling after excluding cortical voxels that can be predicted by features of noninterest (i.e., visual, motor, orthography, and cognitive load; see Materials and Methods for detail). We calculated intramodal predictions using a leave-one-run-out method within the training dataset. Voxels that were consistently predicted in either the MW or ME condition were excluded from

FI analysis; we found significant FI values in the bilateral IPS even after excluding these voxels (15.4% ± 7.2% of voxels were significant; **Figure S3**), indicating that the FI values in the abovementioned analyses were not explained by the effect of sensorimotor processing or general cognitive load.

### Similarities in brain representations related to different mathematical problems

To investigate representational differences between different mathematical problems, we performed RSA based on the encoding model weights. In each anatomical ROI, we calculated a representational similarity matrix (RSM) for both MW and ME problems using Pearson’s correlation distance (**Figure 4A**). We first confirmed similar correlation patterns across subjects for MW (left IPS, *ρ* = 0.436 ± 0.244, Wilcoxon signed-rank test, *p* < 0.001; right IPS, *ρ* = 0.541 ± 0.187, *p* < 0.001) and ME problems (left IPS, *ρ* = 0.491 ± 0.198, *p* < 0.001; right IPS, *ρ* = 0.310 ± 0.171, *p* < 0.001). We thereby compared the resultant RSMs between the two formats and found significant correlations between two RSMs in the bilateral parietal and occipital cortices (**Figure 4B**). In particular, we found a positive correlation between the representational similarities of MW and ME problems in the bilateral IPS (**Figure 4C, D**). The significant correlation existed in the left IPS even after excluding the voxels predicted under the control RW condition (**Figure S4**), indicating that the similarity of the representational relationships among different operators provide the basis of format invariance.

**Figure 4.**
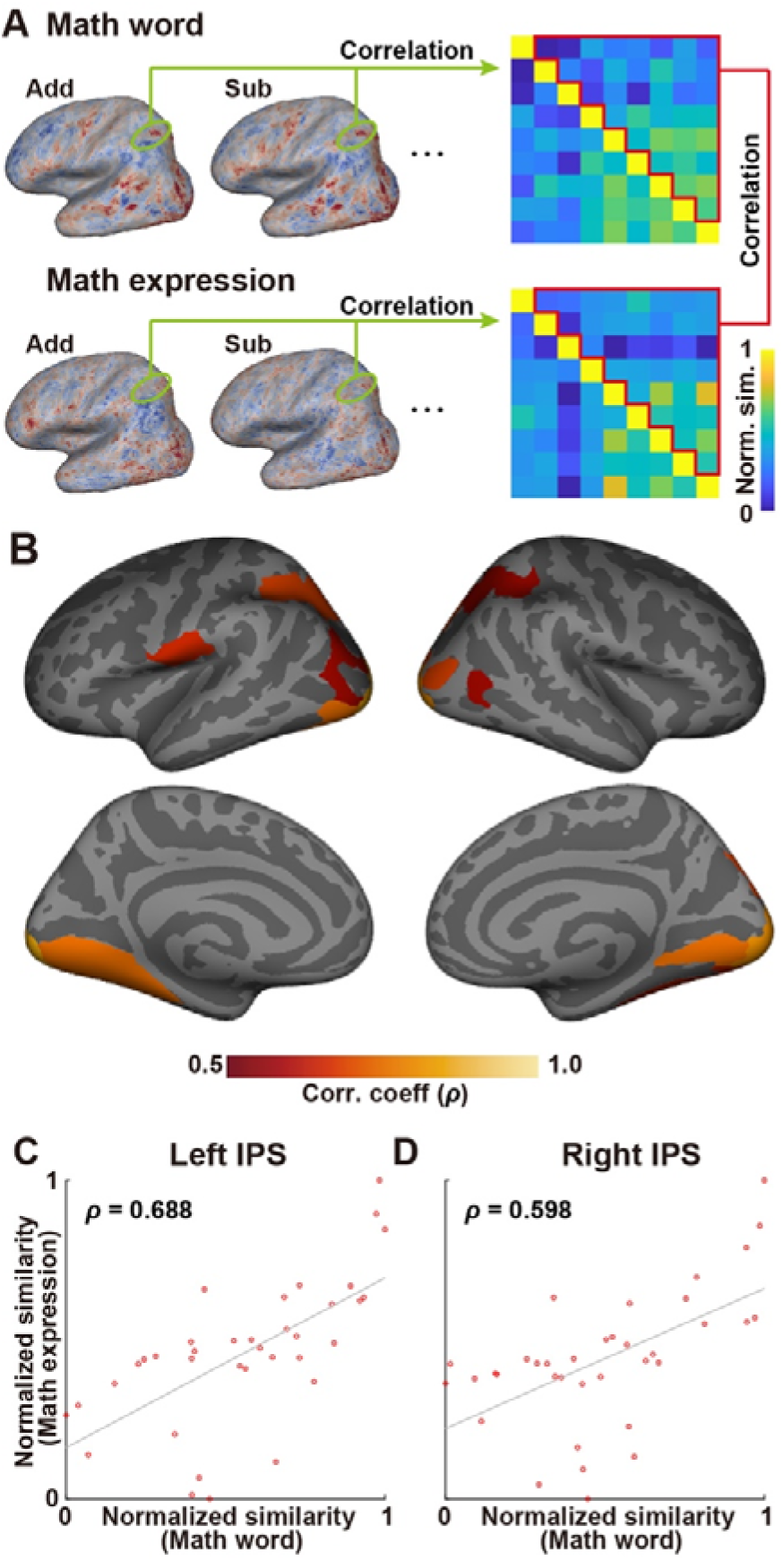
Representational similarity analysis. (**A**) A representational similarity matrix (RSM) was calculated using correlation distances between weight vectors extracted from the target region of interest (ROI) concatenated across all subjects. Spearman’s correlation analysis including the upper triangular parts of the RSM in the math word (MW) and math expression (ME) conditions was conducted for each target ROI. (**B**) Correlation coefficients between RSMs in the MW and ME conditions calculated in each ROI, mapped onto the template brain. Only ROIs with a significant correlation coefficient are shown (*P* < 0.05 with Bonferroni correction). (**C, D**) Scatter plot of the normalized similarities under the MW and ME conditions in the (**C**) left and (**D**) right intraparietal sulcus (IPS). Dots represent elements in the RSMs (surrounded by red lines in **A**).

### Visualization of brain representations related to different mathematical problems

Both FI analysis and RSA revealed shared brain representations for MW and ME problems. To visualize the related organizations, we applied PCA to the weight matrices of encoding models. PCA was also based on the whole cortical voxels. We found that nine tested problems had similar organizations in the MW and ME conditions (**Figure 5A, B**). The representational map visualized for the first and second PCs (PC1 and PC2) showed that double-operator problems were located in the bottom-right area, whereas single-operator problems were located in the upper-left area. Such organization was common across the MW and ME formats, and the mathematical problems were similarly organized even when we used only the bilateral IPS voxels (**Figure S5**). To evaluate the similarity between the representational space of MW and ME problems in a quantitative manner, we calculated Spearman’s correlation coefficients between the PCA loading vectors of two formats. For the top two PCs, the PCA loading vectors were positively correlated between MW and ME problems (PC1 and PC2 were significant using a permutation test, *P* < 0.05; **Figure 5C**). This effect was also observed using the bilateral IPS voxels (PC1 and PC2 were significant using a permutation test, *P* < 0.05; **Figure 5D, S5**). Thus, the nine tested mathematical problems were organized in similar brain representational spaces regardless of their format.

**Figure 5.**
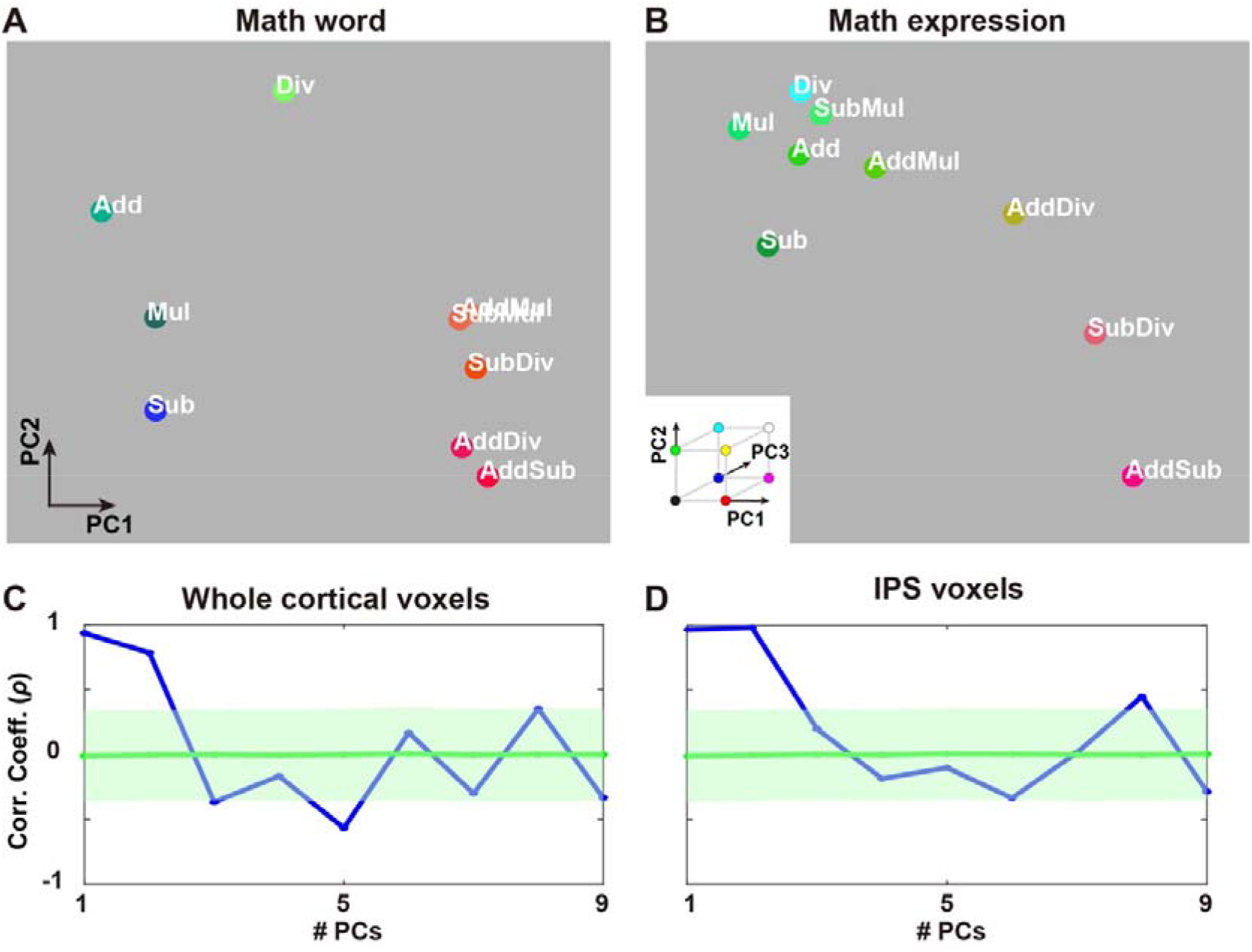
Principal component analysis. (**A, B**) Visualization of the representational space of mathematical problems in the MW (**A**) and ME (**B**) conditions. Colors indicate the loadings of the top three principal components (PC1, red; PC2, green; PC3, blue) of the operator feature weights (concatenated across subjects), which are mapped onto the two-dimensional cognitive space based on the loadings of PC1 and PC2. All operation types are shown in white. (**C, D**) Spearman’s correlation coefficients from analyses of the PC loadings in the MW and ME conditions (blue line) plotted for the top nine PCs using all cortical voxels (**C**) and bilateral IPS voxels (**D**). Mean correction coefficients (cyan line) and standard deviations (shaded areas) were obtained using 5,000 permutations of the operator labels of weight matrices.

## Discussion

To the best of our knowledge, all previous neuroimaging studies on mathematical problem solving have used univariate analysis to assess the format-independency of mathematical problems (Sohn *et al*., 2004; Newman *et al*., 2011; Zhou *et al*., 2018; Chang *et al*., 2019). Thus, it had not been made clear whether the activation overlap of mathematical problems with different formats reflected shared cortical representations. In contrast to univariate analysis, multivariate analysis (i.e., RSA) can test whether activation patterns are similar across different formats. Notably, training and test datasets had not previously been analyzed separately nor had the generalizability of quantitative models to new data been evaluated. In the current study, we demonstrated for the first time that multivariate analyses and encoding model approaches are useful for studying the format-invariance of mathematical problems.

The bilateral IPS is involved in numerical cognition (Eger *et al*., 2003; Cohen Kadosh *et al*., 2005; Cantlon *et al*., 2006; Piazza *et al*., 2007) and processing complex mathematical problems (Nakai & Sakai, 2014; Nakai & Okanoya, 2020). Although format-invariance of quantity information has been questioned by several researchers (Bulthé *et al*., 2014, 2015; Lyons *et al*., 2015; Lyons & Beilock, 2018), we provide evidence that complex mathematical problems are processed in this region in a format-invariant manner. In particular, the left-hemisphere dominance of the IPS in the current study might reflect the specific nature of format-invariance in the case of such complex problems. Note that our results do not indicate format-invariance of symbolic and non-symbolic numbers, as both MW and ME problems used in the current study adopted symbolic numbers based on previous studies (Sohn *et al*., 2004; Zhou *et al*., 2018; Chang *et al*., 2019; Ng *et al*., 2021). Without controlling for the RW condition, we also found FI in the bilateral inferior frontal gyrus (IFG) (**Figure 3A**). The involvement of the left IFG in mathematical problems has been reported in previous studies (Nakai & Sakai, 2014; Nakai *et al*., 2017; Nakai & Okanoya, 2018, 2020), and the functional and anatomical connectivity between the IFG and IPS may play an essential role in the development of mathematical ability (Tsang *et al*., 2009; Emerson & Cantlon, 2012; Rosenberg-Lee *et al*., 2015; Chang *et al*., 2019). Thus, our findings on the IFG and IPS are consistent with the results of other mathematical cognition studies.

Our PCA visualization showed that single- and double-operator problems were distinctly clustered, which is consistent with the result in a previous study comparing single- and double-operator problems (Prabhakaran *et al*., 2001). Researchers have argued that activations in the bilateral IPS during mathematical problem solving are modulated by problem difficulty as measured using reaction times (RTs) (Chang *et al*., 2019). Although our PCA results seemed to reflect a difference between single- and double-operator problems (**Figure S5**), the representational space we obtained after controlling for the RW condition also had a similar organization between single- and double-operator problems (**Figure S4**). In particular, certain double-operator problems were closely located based on the shared single-operator, e.g., the AddDiv and SubDiv conditions and the AddMul and SubMul conditions were paired. Our results for operator-specific representations are consistent with previous reports of double dissociation between different operators (Dehaene & Cohen, 1997) and operator-selective neurons (Kutter *et al*., 2022), which would contribute to the format-invariant predictability of cortical activations.

This study also revealed format-specific aspects of the MW and ME problems. The MW condition is based on natural language sentences and might require the subjects more visual attention and cognitive load than the ME condition. The format-specificity analysis visibly reflected such an influence (Figure 3C, D), indicating a more relevant contribution of the bilateral frontal, temporal, and occipital regions for the MW condition than for the ME condition. In contrast, it is important that the format-invariance was demonstrated in the IPS despite the seemingly large difference between the two conditions. Our method successfully revealed the latent structure of different operators behind the superficial form in which the mathematical problems were presented.

While the overall representations of math operations are similar across the subjects, we also observed notable individual variability. For instance, subjects ID03 and ID06 exhibited relatively smaller MI values than the other subjects (Figure S1). This might be caused by their lower signal-to-noise ratios (Table S1), as their percentage of significantly predicted voxels was smaller than that of the other subjects even for intra-format analyses. There might be room for improving the MI value definition in that it is affected by intra-format prediction accuracy values.

Although the current study paves the way toward quantitative modeling of complex mathematical problems, we acknowledge that it has several limitations. First, because we chose experimental stimuli from existing datasets (Roy & Roth, 2015; Roy *et al*., 2015), we could not perfectly balance the number of letters and digit sizes. Although we excluded such effects by subtracting baseline data determined using the RW condition with the same visual stimuli (**Figure 3B**) and by explicitly regressing out features of noninterest (**Figure S3**), the between-condition difference may still have had some influence on the representational organization of the tested math problems. Second, although we validated our results for each subject (**Figure S2**) and the test sample size (N = 465 for the ME and N = 605 for the MW conditions), as well as the number of subjects, were consistent with those of previous studies (Kay, Naselaris, *et al*., 2008; Nishimoto *et al*., 2011; Huth *et al*., 2016; Horikawa *et al*., 2020; Nakai & Nishimoto, 2020), the relatively small number of subjects tested may limit the generalizability of our results to the broader population and other age groups. Third, the categorical model of mathematical operations (i.e., operator features) may not fully capture the complex processes involved in math problem solving, especially the quantity information used in the problems. Therefore, further studies are warranted to fully clarify the individual differences and detailed subprocesses involved in mathematical problems. Nevertheless, based on the current work, we conclude that the bilateral IPS network subserves format-invariant representations of mathematical problems.

## Materials and Methods

### Subjects

Eight healthy college students (aged 20–23 years, three females, all with normal vision), denoted as ID01–ID08, participated in this study. The number of subjects is in the same range as previous encoding model studies (Kay, Naselaris, *et al*., 2008; Nishimoto *et al*., 2011; Huth *et al*., 2016; Horikawa *et al*., 2020; Nakai & Nishimoto, 2020). The small number of subjects is compensated for by a large number of samples for each subject (i.e., 3 h of tests). Subjects were all right-handed (laterality quotient = 80–100) as assessed using the Edinburgh inventory (Oldfield, 1971). Written informed consent was obtained from all subjects prior to their participation in the study, and the experiment was approved by the ethics and safety committee of the National Institute of Information and Communications Technology in Osaka, Japan.

### Stimuli and testing procedure

Subjects performed arithmetic problems in two different formats, namely MW and ME problems (see **Figure S6** for the behavioral results). We selected MW problems with a single operation of addition (Add), subtraction (Sub), multiplication (Mul), and division (Div) from the IL dataset (Roy & Roth, 2015), and MW problems with two operations, including addition and subtraction (AddSub), addition and multiplication (AddMul), addition and division (AddDiv), subtraction and multiplication (SubMul), and subtraction and division (SubDiv), from the CC dataset (Roy *et al*., 2015). Each condition consisted of 35 instances. Both of the abovementioned datasets have been widely used in previous machine learning studies on MW problems (Mandal & Naskar, 2019). For our MW problems, the original English sentences were translated into Japanese by the first author (T.N.). Corresponding arithmetic expression problems were created based on the MW problems; thus, these problems gave the same results.

In the MW condition, an arithmetic word problem, (e.g., “There are 3 eggs in each box. How many eggs are in 2 boxes?”) was presented for 6 or 10 s [6 s for the single-operator conditions (Add, Sub, Mul, and Div) and 10 s for the double-operator conditions (AddSub, AddMul, AddDiv, SubMul, and SubDiv)]. Subjects had access to a two-button response pad in their left hand, and they were instructed to perform a calculation based on the presented problem and press the left button when they had calculated an answer. After 1–2 s during which a fixation cross stimulus was displayed, a probe digit stimulus was presented for 2 s (e.g., “6”). Subjects were asked to press the left button if the presented digit matched their answer and the right button if it did not match their answer. The next trial started after 1–2 s, during which time a fixation cross stimulus was shown. The number of letters used in each condition was as follows: 61.3 ± 13.3 (Add), 57.0 ± 6.6 (Sub), 51.4 ± 11.2 (Mul), 66.9 ± 12.0 (Div), 89.4 ± 13.4 (AddSub), 85.6 ± 13.1 (AddMul), 91.2 ± 16.3 (AddDiv), 83.8 ± 7.2 (SubMul), and 91.4 ± 13.6 (SubDiv).

In the ME condition, an arithmetic expression problem (e.g., “3 × 2 = ?”) was presented for 4 or 6 s [4 s for the single-operator conditions (Add, Sub, Mul, and Div), and 6 s for the double-operator conditions (AddSub, AddMul, AddDiv, SubMul, and SubDiv)]. As in the MW condition, the subjects were instructed to perform a calculation based on the presented problem and to press the left button when they had calculated an answer. The fixation cross stimulus was then presented for 1–2 s, after which a probe digit stimulus was presented for 2 s (e.g., “6”). Subjects were again asked to press the left or right button if the presented digit matched or did not match their answer, respectively. The next trial began after the fixation cross stimuli was again displayed for 1–2 s. The number of letters used in each condition was as follows: 5.5 ± 0.5 (Add), 6.5 ± 0.5 (Sub), 5.0 ± 0.2 (Mul), 5.9 ± 0.4 (Div), 9.3 ± 0.9 (AddSub), 8.9 ± 0.3 (AddMul), 10.1 ± 0.7 (AddDiv), 9.6 ± 0.5 (SubMul), and 10.9 ± 0.5 (SubDiv).

In the RW condition, an arithmetic word problem was presented for 6 or 10 s as described above for the MW condition. The set of sentences presented was the same as that used in the MW condition (e.g., “There are 3 eggs in each box. How many eggs are in 2 boxes?”). However, subjects were instructed to carefully read the sentence and press the left button when they had memorized all non-numerical nouns. After a fixation cross stimulus was shown for 1–2 s, two words were presented for 2 s (e.g., “eggs” and “boxes”). Subjects were asked to press the left button if both nouns were included in the original sentence and the right button if one of the two nouns were not included in the sentence. Two nouns were presented in the answer-matching phase to ensure that the subjects memorized all the nouns that appeared in the sentence. The next trial started after a fixation cross stimuli was displayed for 1–2 s.

Stimuli were presented on a projector screen inside the scanner (21.0 × 15.8° visual angle at 30 Hz). During scanning, subjects wore MR-compatible ear tips. Presentation software (Neurobehavioral Systems, Albany, CA, USA) was used to control the stimulus presentation and collection of behavioral data. To measure button responses, optic response pads with two buttons were used (HHSC-2×2, Current Designs, Philadelphia, PA, USA).

The experiment was performed for three days, with six runs performed each day. In total, 16 runs were conducted, of which 6 training and 2 test runs were in the MW condition, and another 6 training and 2 test runs were in the ME condition. The presentation order was as follows: day 1, MW, ME, MW, ME, MW, and RW (57.5 min); day 2, MW, ME, MW, ME, MW, and ME (54.5 min); day 3, RW, ME, MW, ME, MW, and ME (54.5 min). A T1 anatomical image (6 min) was acquired at the end of the first day. The presentation order was identical for all subjects. The data from each pair of test runs were averaged to increase the signal-to-noise ratio. Each run contained 90 trials. The duration of a single run was 595 s for the MW condition and 455 s for the ME condition. At the beginning of each run, 10 s of dummy scans were acquired, during which the fixation cross was displayed, and these dummy scans were later omitted from the final analysis to reduce noise. We also obtained 10 s of scans at the end of each run, during which the fixation cross was displayed; however, these scans were included in the analyses.

### MRI data acquisition

The experiment was conducted using a 3.0 T scanner (MAGNETOM Prisma; Siemens, Erlangen, Germany) with a 64-channel head coil. We scanned 72 interleaved axial slices that were 2-mm thick without a gap, parallel to the anterior and posterior commissure line, using a T2*-weighted gradient echo multiband echo-planar imaging sequence [repetition time (TR) = 1,000 ms; echo time (TE) = 30 ms; flip angle (FA) = 62°; field of view (FOV) = 192 × 192 mm^2^; resolution = 2 × 2 mm^2^; multiband factor = 6]. We obtained 605 volumes for the MW and RW conditions and 465 volumes for the ME condition, with each set following 10 dummy images. For anatomical reference, high-resolution T1-weighted images of the whole brain were also acquired from all subjects with a magnetization-prepared rapid acquisition gradient echo sequence (TR = 2,530 ms; TE = 3.26 ms; FA = 9°; FOV = 256 × 256 mm^2^; voxel size = 1 × 1 × 1 mm^3^).

### fMRI data preprocessing

Motion correction in each run was performed using the statistical parametric mapping toolbox (SPM12; Wellcome Trust Centre for Neuroimaging, London, UK; http://www.fil.ion.ucl.ac.uk/spm/). All volumes were aligned to the first EPI image for each subject. Low-frequency drift was removed using a median filter with a 120-s window. Slice timing correction was performed against the first slice of each scan. The response for each voxel was then normalized by subtracting the mean response and scaling it to the unit variance. We used FreeSurfer (https://surfer.nmr.mgh.harvard.edu/) to identify the cortical surfaces from the anatomical data and to register these to the voxels of the functional data. For each subject, the voxels identified in the cerebral cortex (59,499–75,980 voxels per subject) were used in the analysis.

### Operator features

The operator features were composed of one-hot vectors, which were assigned values of 1 or 0 for each time bin during the stimulus presentation of arithmetic problems, indicating whether one of the nine tested conditions (Add, Sub, Mul, Div, AddSub, AddMul, AddDiv, SubMul, and SubDiv) was performed in that period; thus, nine operator features were used.

### Motion energy features

We employed a motion energy (MoE) model used in previous studies (Nishimoto *et al*., 2011; Koide-Majima *et al*., 2020; Nakai & Nishimoto, 2020) that can be found in a public repository (https://github.com/gallantlab/motion_energy_matlab). First, movie frames and pictures were spatially downsampled to 96 × 96 pixels. The RGB pixel values were then converted into the Commission International de l’Eclairage LAB color space, and the color information was discarded. The luminance (L*) pattern was passed through a bank of three-dimensional spatiotemporal Gabor wavelet filters; the outputs of the two filters with orthogonal phases (quadrature pairs) were squared and summed to yield local MoE. Subsequently, MoE was compressed with a log-transformation and temporally downsampled to 0.5 Hz. Filters were tuned to six spatial frequencies (0, 1.5, 3.0, 6.0, 12.0, and 24.0 cycles per image) and three temporal frequencies (0, 4.0, and 8.0 Hz) without directional parameters. Filters were positioned on a square grid that covered the screen. The adjacent filters were separated by 3.5 standard deviations of their spatial Gaussian envelopes. To reduce the computational load, the original MoE features, which had 1,395 dimensions, were reduced to 300 dimensions using principal component analysis.

### Button response feature

The button response (BR) feature was constructed based on the number of button responses per second. One BR feature was used.

### Letter feature

The letter feature was constructed based on the number of letters that appeared in each stimulus. One letter feature was used.

### Reaction time feature

For each time bin during the presentation of a mathematical problem, the reaction time (RT) of that trial was assigned as a cognitive load of the trial. One RT feature was used.

### Accuracy feature

For each time bin during the presentation of a mathematical problem, the accuracy of the operator type of the given problem (for the given subject) was assigned as another cognitive load index. One accuracy feature was used.

### Encoding model fitting

In the encoding model, the cortical activity in each voxel was fitted with a finite impulse response model that captured the slow hemodynamic response and its coupling with neural activity (Kay, David, *et al*., 2008; Nishimoto *et al*., 2011). The feature matrix F_E_ [T × 6N] was modeled by concatenating sets of [T × N] feature matrices with six temporal delays of 2–7 s (T = number of samples; N = number of features). The cortical response R_E_ [T × V] was then modeled by multiplying the feature matrix F_E_ by the weight matrix W_E_ [6N × V] (V = number of voxels):

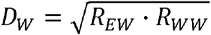

We used an L2-regularized linear regression with the training dataset to obtain the weight matrix W_E_. The training dataset consisted of 3,630 (3,630 s) and 2,790 (2,790 s) samples for the MW and ME conditions, respectively, whereas a training dataset was not provided for the RW condition. The optimal regularization parameter was assessed using 10-fold cross-validation where the 11 different regularization parameters ranged from 1 to 2^10^.

The test dataset consisted of 605 samples (605 s) for the MW and RW conditions and 465 samples (465 s) for the ME condition (each repeated twice). Two repetitions of the test dataset were averaged to increase the signal-to-noise ratio. Statistical significance (one-sided) was computed by comparing estimated correlations to the null distribution of correlations between two independent Gaussian random vectors with the same length as the test dataset. The statistical threshold was set at *P* < 0.05 and corrected for multiple comparisons using the FDR procedure (Benjamini & Hochberg, 1995). For data visualization on the cortical maps, we used pycortex (Gao *et al*., 2015) and fsbrain (Schäfer & Ecker, 2020).

### Encoding model fitting excluding regressors of noninterest

To evaluate the possible effect of sensorimotor features and general cognitive load on the model predictions, we performed an additional encoding model fitting by excluding the features of noninterest. To this end, we concatenated the MoE (visual), BR (motor), RT (cognitive load), accuracy (cognitive load), and letter (orthographic) features. The concatenated features were used as a feature matrix for the encoding modeling. The original training dataset was divided into five subtraining runs and a single subtest run. Encoding models were trained using the subtraining dataset, and prediction accuracy was calculated using the remaining subtest dataset. L2-regularized linear regressions were applied in the same manner as that described in the “Encoding model fitting”subsection in the Methods. This procedure was repeated for all runs in the original training dataset. Cortical voxels significantly predicted in at least five out of six repetitions were regarded as nonmath voxels and excluded from the analysis of target encoding models.

### Quantification of format invariance

To quantify how the unimodal models explained brain activity in each voxel regardless of the presentation format, we defined FI values. In our previous study, we used a similar measurement to quantify how linguistic information is processed in a modality-invariant manner (Nakai, Yamaguchi, *et al*., 2021). Note that the variance partitioning analysis (de Heer *et al*., 2017) cannot be used to analyze the format-invariance in the current experiment because this analysis compares explained variances of two (or more) different types of features extracted from the same dataset. In contrast, this experiment aims at comparing the same operator features extracted from two datasets. To quantify FI based on prediction accuracy, we used the geometric mean of prediction accuracy rather than the weight correlation. This was justified because models with similar weight values have similar predictive performance. FI consisted of two components: D_W_ and D_E_. D_W_ was defined as the degree of predictability for the MW test dataset regardless of the training format: where R_WW_ and R_EW_ are the intramodal prediction accuracy for the MW model and the cross-modal prediction accuracy for the ME model when applied to the test dataset for the MW condition, respectively. Similarly, D_E_ was defined as the degree of predictability calculated for the ME test dataset regardless of the training format:

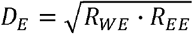

where R_EE_ and R_WE_ are the intra-format prediction accuracy using an encoding model trained with the ME condition and the cross-format prediction accuracy trained with the MW condition when applied to the test dataset for the ME condition, respectively. For all voxels with negative prediction accuracies, the prediction accuracy was reset to 0 to avoid obtaining imaginary values. FI was then calculated for each voxel as a geometric mean between *D_w_* and *D_E_* as follows:

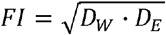

FI values ranged from 0 to 1; high FI indicates that the target features are represented in a format-invariant manner, whereas an FI of zero indicates that the target voxel does not have a shared representation for MW and ME problems. To calculate the null distribution of FI values, a set of four correlation coefficients produced using Gaussian random vector pairs were used instead of R_WW_, R_EE_, R_EW_, and R_WE_. The statistical threshold was set at *P* < 0.05 and corrected for multiple comparisons using the FDR procedure (Benjamini & Hochberg, 1995).

### Format specificity

To quantify how the encoding models explained brain activity that was specific for a certain format, we defined format specificity, which was calculated in each voxel for each format (FS_W_ for the MW problems and FS_E_ for the ME problems) as the difference between the intra-format and cross-format prediction accuracies:

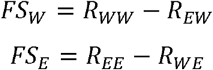

FS value ranges from −1 to 1. A high FS value indicates that the target features are represented specifically according to the target format, where negative FS indicates that the target voxel does not have a format-specific representation. Significance and FDR corrections for multiple comparisons were calculated as described for the FI values.

### RSA

For each of the target anatomical ROIs, *operator similarity* was calculated using the Pearson correlation distance between weight vectors of different operators extracted from the weight matrix of operator feature encoding models. We selected the voxels that showed significant FI values and averaged six time delays for each operator. An RSM was calculated based on all combinations of nine operator types, using concatenated feature vectors across all subjects. The upper triangular part of the RSM was rearranged into a single vector for both the MW and ME conditions, and Spearman’s correlation coefficients for correlations between the two rearranged vectors were calculated.

### PCA

For each subject, we performed PCA on the weight matrix of the operator feature model concatenated across the eight subjects. We selected the voxels that showed significant FI values and averaged six time delays for each operator. We applied PCA to the [9*V] weight matrices (V = number of voxels) of the MW and ME encoding models. To show the structure of the representational space of mathematical problems, nine operators were mapped onto the two-dimensional space using the loadings of the first and second PCs, i.e., PC1 and PC2, as the x-axis and y-axis, respectively. In this space, the operators were colored red, green, and blue based on the relative PCA loadings in PC1, PC2, and PC3, respectively. To investigate the relationship between the components obtained using the MW and ME models, we calculated Spearman’s correlation coefficients for correlations between the PCA scores for each component of two models (e.g., between PC1 of the MW model and PC1 of the ME model). To obtain a null distribution of such correlation coefficients, we randomly permutated the categorical labels of the MW and ME models 5,000 times before applying PCA. *P* values of the actual correlation coefficients were evaluated according to the number of random correlation coefficients larger than the actual value across all permutations.

## Data and code availability

The source data and analysis code used in the current study are available from Zenodo (https://doi.org/10.5281/zenodo.6605258).

## Acknowledgments

We thank MEXT/JSPS KAKENHI [grant numbers JP17K13083 and JP18H05091 in #4903 (Evolinguistics) for T.N., and JP15H05311 and JP18H05522 for S.N.] as well as JST CREST (JPMJCR18A5) and ERATO (JPMJER1801) (for S.N.) for their partial financial support of this study. The funders played no role in the study design, data collection, and analysis, decision to publish, or preparation of the manuscript.

## Author contributions

T.N. and S.N. designed the study; T.N. collected and analyzed the data; T.N. and S.N. wrote the manuscript.

## Competing interests

The authors declare no competing interests.

## Abbreviation

MW: math word
ME: math expression
RW: read word
IPS: intraparietal sulcus
IFG: inferior frontal gyrus
RSA: representational similarity analysis
RSM: representational similarity matrix
PCA: principal component analysis
fMRI: functional magnetic resonance imaging
FI: format invariance
FS: format specificity
ROI: region of interest
RT: reaction time
TR: repetition time
TE: echo time FA flip angle
FOV: field of view
MoE: motion energy
BR: button response

## Supplementary Information

**Figure S1.**
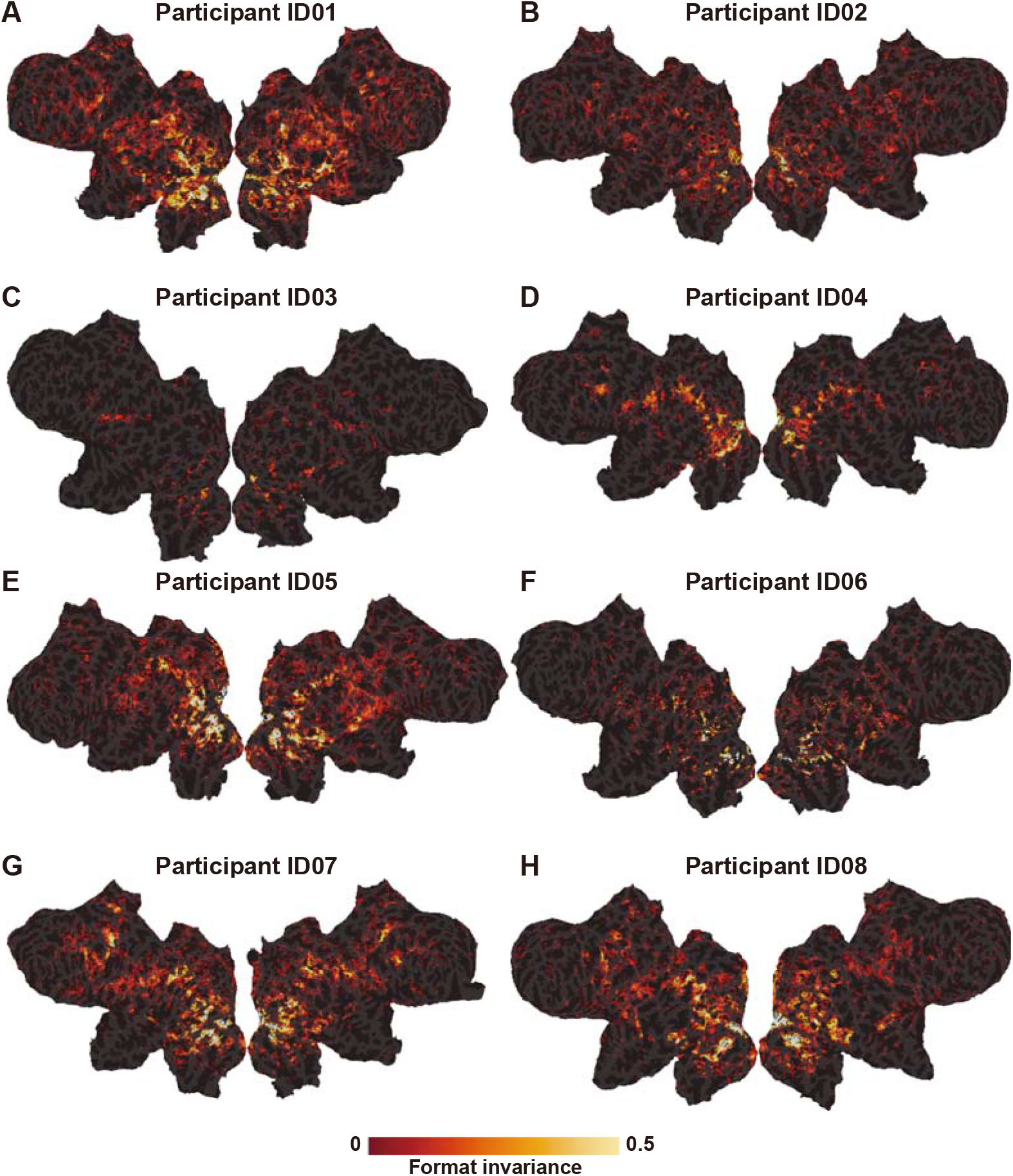
Format invariance map for each subject. Cortical maps of format invariance for subjects ID01–ID08 mapped onto cortical surfaces. Only significant voxels (*P* < 0.05, FDR corrected) are shown.

**Figure S2.**
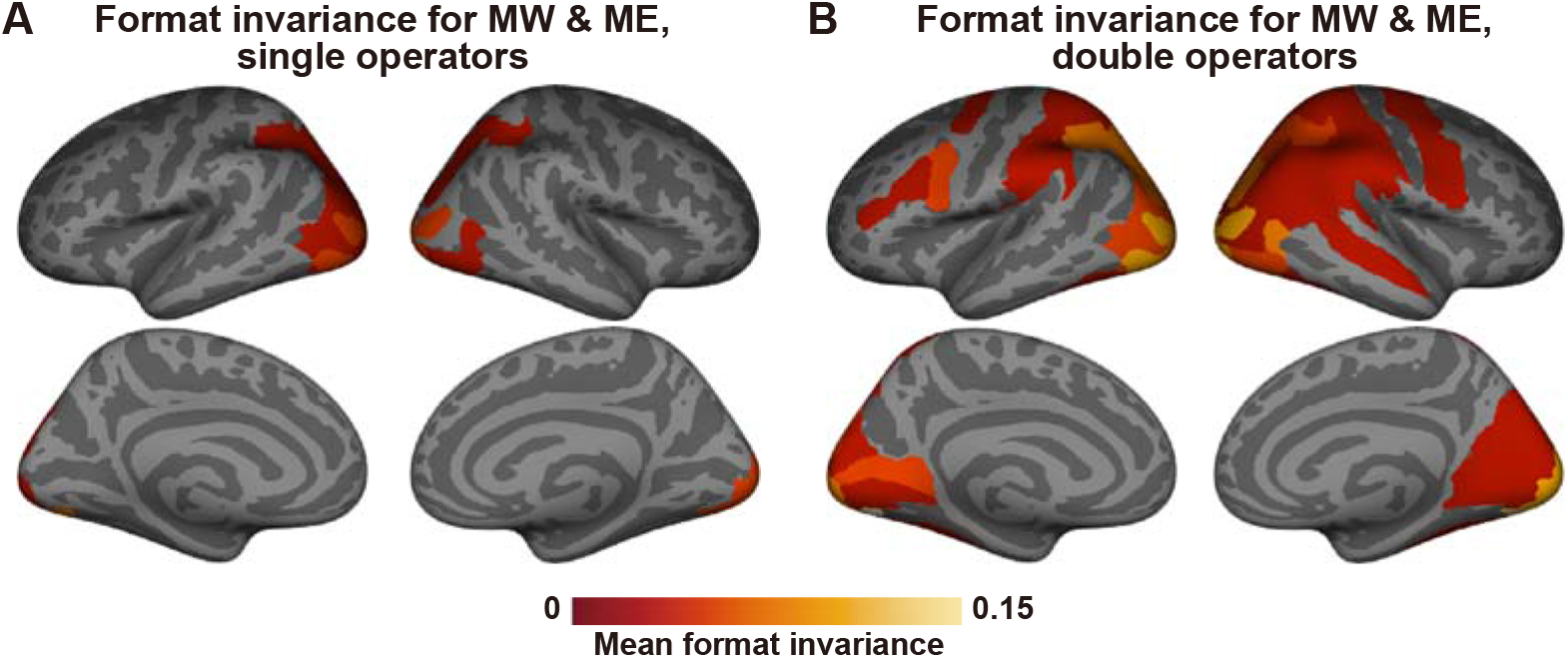
Format invariance analysis of single- and double-operators. Group averages of format invariance (FI) for (**A**) single- and (**B**) double-operator problems mapped onto the template brain. FI values were averaged in each anatomical region of interest (ROI) and averaged across eight subjects. Only ROIs with >20% significant voxels are shown. MW, math word; ME, math expression.

**Figure S3.**
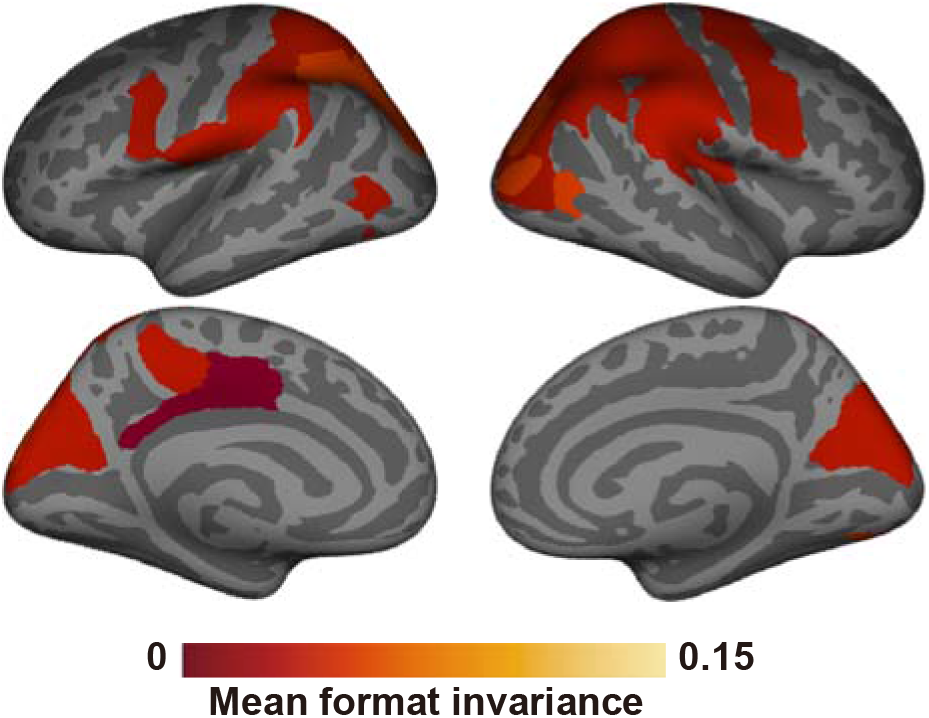
Format invariance analysis after excluding the effects of regressors of noninterest. Group averages of format invariance after excluding voxels significantly predicted by regressors of noninterest, either in the math word or math expression problems. The regressors of noninterest consisted of the visual, motor, orthographic, and cognitive load features.

**Figure S4.**
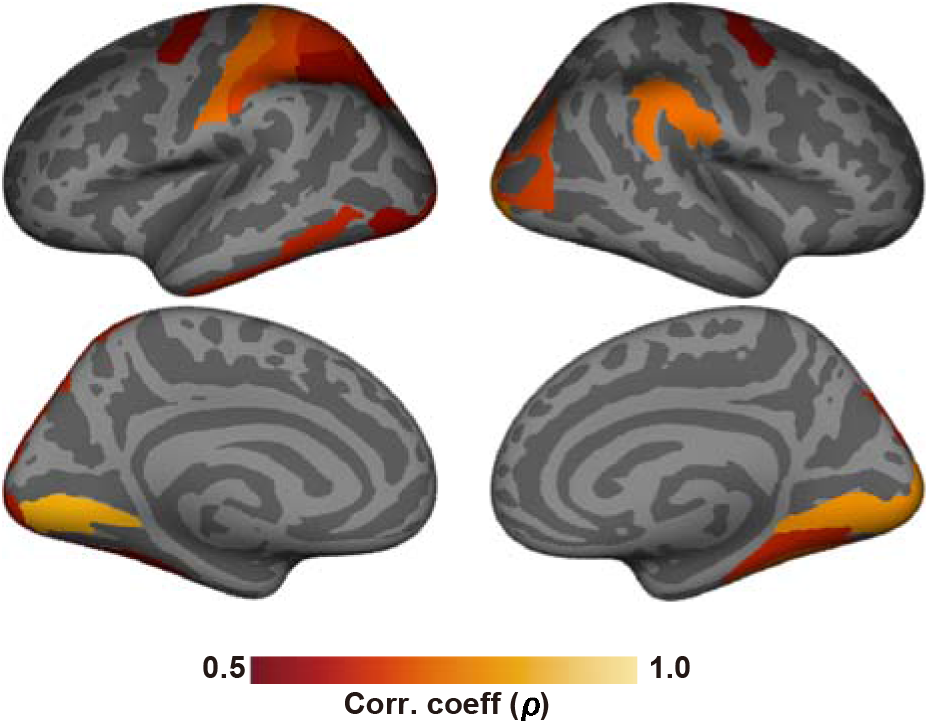
Representational similarity analysis using the control condition. Correlation coefficients between representational similarity matrices in the MW and ME conditions, calculated in each ROI and mapped onto the template brain. Significant voxels after subtraction of the prediction accuracy in the read word condition were used. Only ROIs with a significant correlation coefficient are shown (*P* < 0.05 with Bonferroni correction).

**Figure S5.**
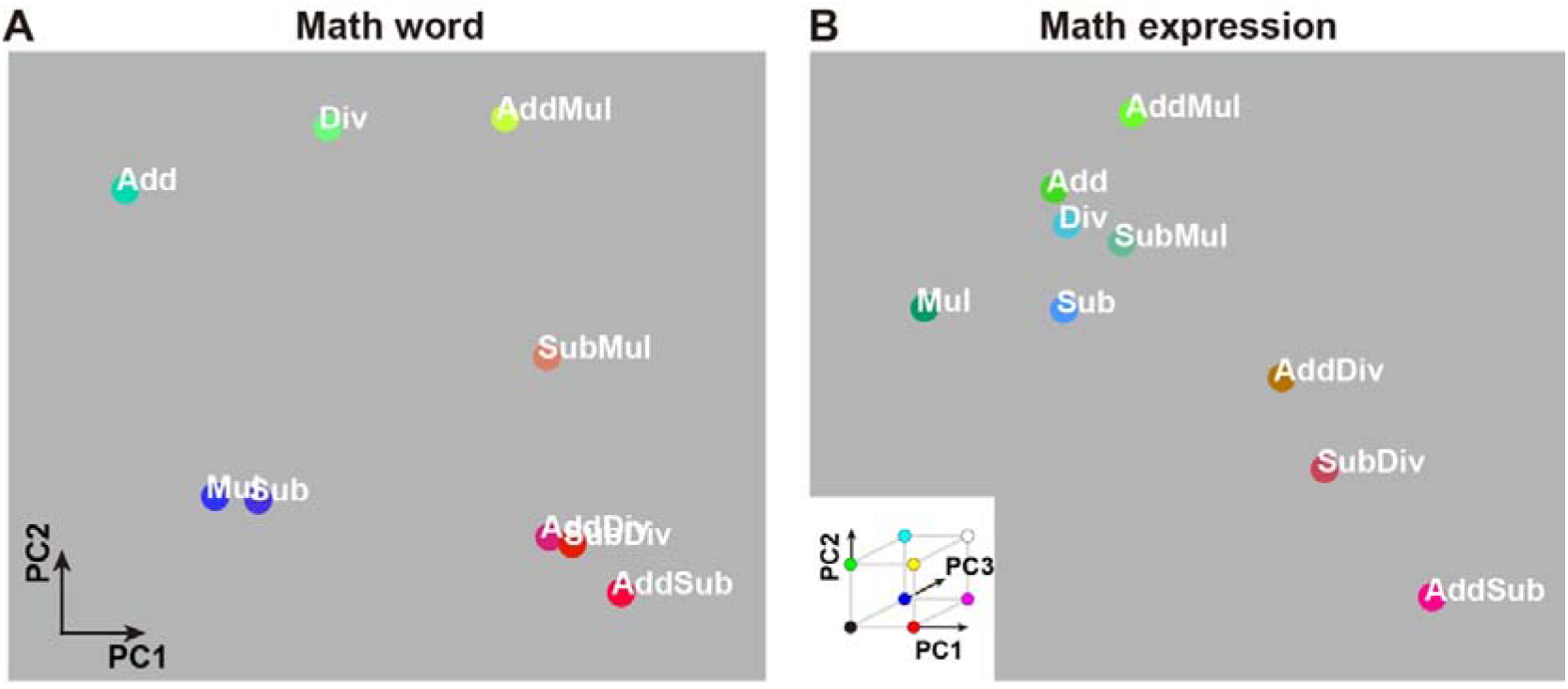
Principal component analysis using the bilateral IPS voxels. (**A, B**) Visualization of the representational space of mathematical problems in (**A**) the math word and (**B**) the math expression conditions using the bilateral IPS voxels. Colors indicate the loadings of the top three principal components (PC1, red; PC2, green; PC3, blue) of the operator feature weights (concatenated across subjects), which were mapped onto the two-dimensional cognitive space based on the loadings of PC1 and PC2.

### Behavioral results

The subjects were able to solve both MW and ME conditions with >90% accuracy (MW condition, 92.7% ± 4.0%; ME condition, 94.8% ± 2.5%; Cohen’s *d* = 0.62; **Figure S6A** and **S6C**, respectively). The subjects solved the single-operator (Add, Sub, Mul, and Div) and double-operator (AddSub, AddMul, AddDiv, SubMul, and SubDiv) problems with equal accuracy both in the MW (single-operator, 93.0% ± 4.2%; double-operator, 92.5% ± 4.5%; *d* = 0.11) and ME (single-operator, 94.5% ± 3.1%; double-operator, 95.1% ± 2.5%; *d* = 0.22) conditions. These results indicate that subjects did not have specific difficulty in solving arithmetic problems in either format.

As expected, RT was higher in the MW condition than that in the ME condition (MW condition, 4.59 ± 0.84 s; ME condition, 1.56 ± 0.30 s; *d* = 2.70; **Figures S6B** and **S6D**, respectively). For the MW condition, RTs were significantly higher in the double-operator conditions than those in the single-operator conditions for all subjects (single-operator, 3.32 ± 0.63 s; double-operator, 5.60 ± 1.02 s; *d* = 2.70). For the ME condition, RTs were higher in the double-operator problems than those in the single-operator problems (single-operator, 1.06 ± 0.14 s; double-operator, 1.96 ± 0.45 s; *d* = 2.69).

For the control RW condition, there was no distinction among different operators, but there was a distinction between sentences used in the single-operator and double-operator problems. Overall accuracy in the RW condition was comparable to that in the MW condition (89.6% ± 4.5%; *d* = 0.67), whereas the accuracy was lower than that in the ME condition (*d* = 1.37) (**Figure S6E**, left). RT in the RW condition was comparable with that in the MW condition (4.16 ± 0.86 s; *d* = 0.61), but it was higher than that in the ME condition (*d* = 4.02) (**Figure S6E**, right). Additionally, RT was higher in the double-operator sentences (single-operator, 3.41 ± 0.62 s; double-operator, 4.76 ± 1.09 s; *d* = 1.53). In contrast, accuracy was higher in the single-operator sentences (single-operator, 92.5% ± 4.2%; double-operator, 87.3% ± 5.6%; *d* = 1.06). These results indicate that the RW and MW conditions had similar levels of difficulty for the subjects, but that the former was more demanding than the ME condition.

**Figure S6.**
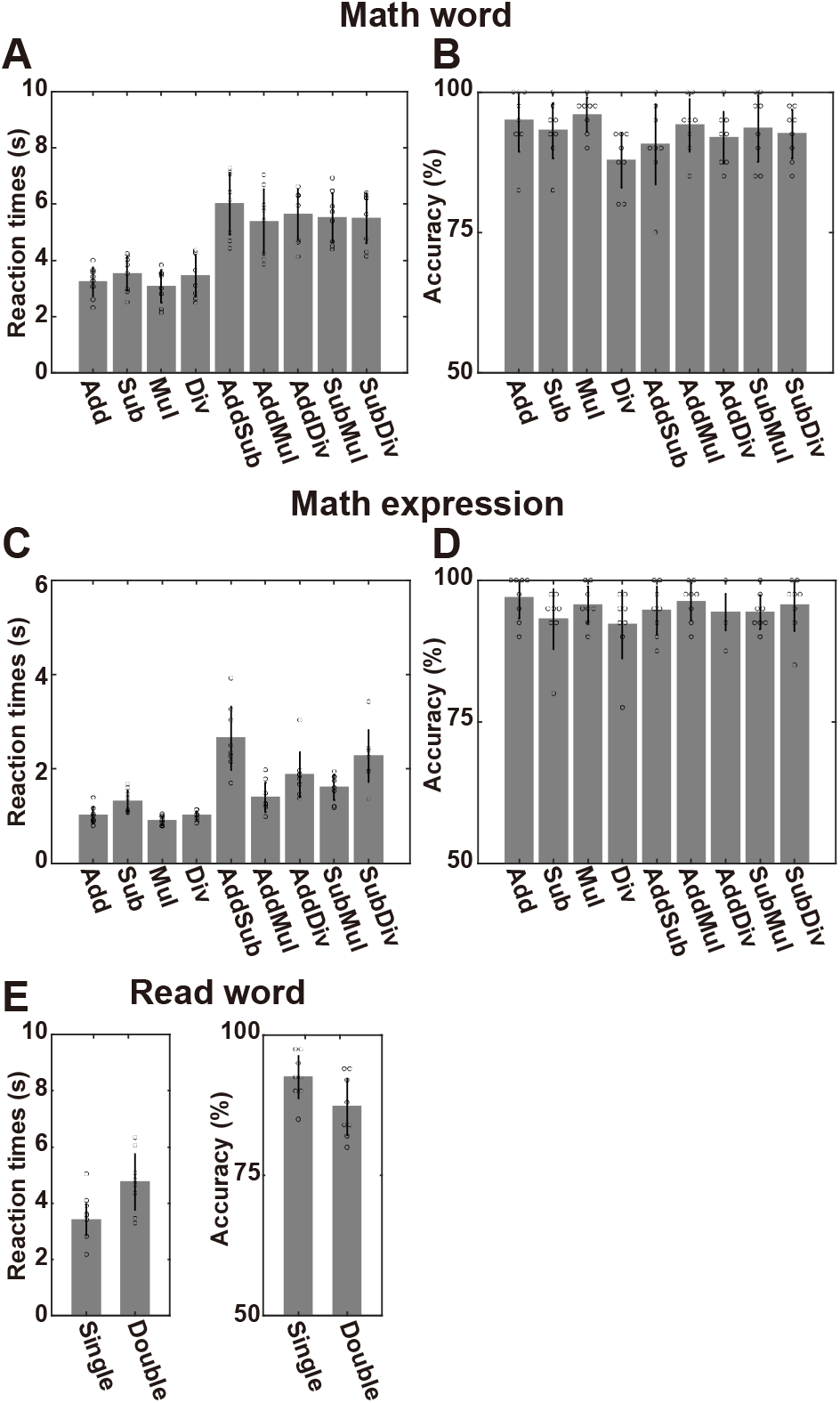
Behavioral results. Accuracy (**A** and **C**) and reaction time (**B** and **D**) of the math word and math expression conditions, respectively, are shown for nine operator types. (**E**) Accuracy and reaction time of the read word condition. Data for each subject are indicated by dots, and error bars represent standard deviation.

**Table S1.**
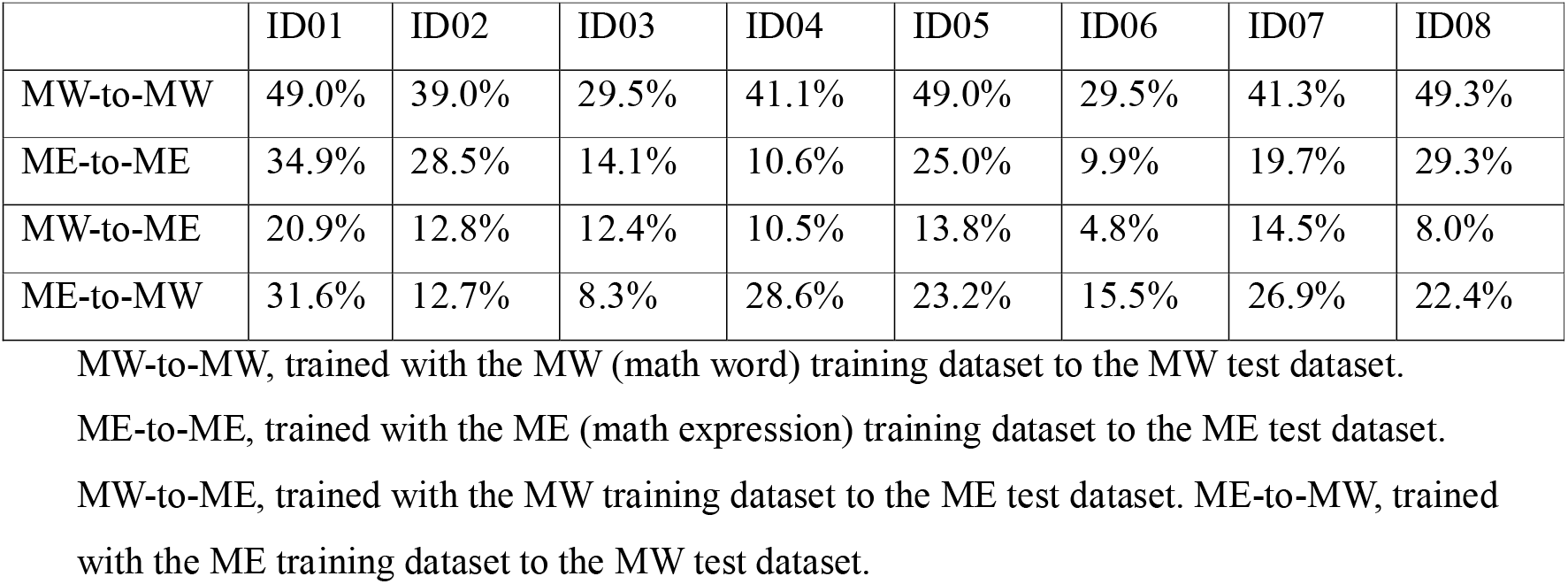
Percentage of significantly predicted voxels across the cortex.

## Notes

### Competing Interest Statement

The authors have declared no competing interest.

### Summary of Updates

Figures 1-4 revised.

